# Colon epithelial cell-specific Bmal1 deletion impairs bone formation in mice

**DOI:** 10.1101/2021.08.02.454190

**Authors:** Frank C. Ko, Sarah B. Jochum, Brittany M. Wilson, Amal Adra, Nikhil Patel, Sherry Wilber, Maliha Shaikh, Christopher Forsyth, Ali Keshavarzian, Garth R. Swanson, D. Rick Sumner

## Abstract

The circadian clock system regulates multiple metabolic processes, including bone metabolism. Previous studies have demonstrated that both central and peripheral circadian signaling regulate skeletal growth and homeostasis. Disruption in central circadian rhythms has been associated with a decline in bone mineral density and the global and osteoblast-specific disruption of clock genes in bone tissue leads to lower bone mass. Gut physiology is highly sensitive to circadian disruption. Since the gut is also known to affect bone remodeling, we sought to test the hypothesis that circadian signaling disruption in colon epithelial cells affects bone. We therefore assessed structural, functional, and cellular properties of bone in 8 week old Ts4-Cre and Ts4-Cre;Bmal1^fl/fl^ (cBmalKO) mice, where the clock gene Bmal1 is deleted in colon epithelial cells. Axial and appendicular trabecular bone volume was significantly lower in cBmalKO compared to Ts4-Cre 8-week old mice in a sex-dependent fashion, with male but not female mice showing the phenotype. Similarly, the whole bone mechanical properties were deteriorated in cBmalKO male mice. The tissue level mechanisms involved suppressed bone formation with normal resorption, as evidenced by serum markers and dynamic histomorphometry. Our studies demonstrate that colon epithelial cell-specific deletion of Bmal1 leads to trabecular and cortical bone loss in male mice.

## Introduction

Circadian rhythms are endogenous 24-hour recurring behavioral, physiological, and metabolic patterns that are orchestrated by the circadian clock (1, 2). Central circadian misalignment occurs when there is a mismatch between an environmental cue (e.g., light) and the central clock in the suprachiasmatic nucleus, while peripheral circadian misalignment occurs when there is a mismatch between an environmental cue (e.g., time of feeding) and endogenous rhythms of a specific organ (e.g., the gut). Disruption in circadian rhythms due to rotating-shift work or long distance travel with jet leg, has been associated with several metabolic disorders, including obesity, insulin resistance, hypertension, and diabetes (3–6). Clinical studies have also demonstrated that skeletal deterioration is associated with disrupted circadian rhythms (7). For example, in studies of sleep deprived individuals, circadian rhythm disruption is associated with decreased bone mineral density, increased fracture risk, and lower serum bone formation markers (8–12). The mechanisms of how the circadian clock system regulates bone metabolism and health remain unclear (13).

Genes such as the brain and muscle ARNT-like protein-1 (Bmal1), circadian locomotor output cycles kaput (Clock), Period1 (Per1), Period2 (Per2), cryptochrome1 (Cry1), and cryptochrome2 (Cry2) are integral to the circadian signaling pathway (14). These genes have also been shown to regulate skeletal growth and bone remodeling. When Bmal1 was globally deleted in mice, longitudinal skeletal growth and osteoblast and osteocyte numbers were decreased (15). Osteoprogenitor and osteoblast-specific deletion of Bmal1 led to decreased bone due to increased bone resorption (16). Global deletion of Per1/2 or Cry1/2, which are negative regulators of circadian signaling, increased bone in mice (17). More recently, intestinal epithelium-specific deletion of Bmal1 was shown to decrease bone in mice (Villin-Cre;Bmal1^fl/fl^) (18). With this model, Bmal1 is deleted in both the small intestine and the large intestine (19) and likely in the kidney as well since Villin-Cre is expressed in kidney proximal tubules (20, 21). The study demonstrated that the bone deficit was driven by increased bone resorption due to vitamin D-induced calcium malabsorption in the duodenum to maintain normocalcemia and suppressed bone formation due to altered sympathetic tone. However, no study to date has examined the effects of colon-specific Bmal1 deletion on bone.

We recently demonstrated that Bmal1 deletion in the colon epithelium does not cause ulcerative colitis but still leads to mild colonic inflammation in mice [S]. Several studies show that alterations in gut physiology, such as inflammatory bowel diseases, have consequences for skeletal health (22–25) and moderate gut inflammation that does not cause weight loss leads to bone loss in mice (26). We therefore sought to determine the effects of Bmal1 deletion in colon epithelium on bone. We hypothesized that colon epithelial cell-specific deletion of Bmal1 will decrease bone in mice.

## Materials and Methods

### Mouse

Animal studies were approved by the Rush University Medical Center Institutional Animal Care and Use Committee. Ts4-Cre (Ts4-Cre) and Ts4-Cre;Bmal1^fl/fl^ (cBmalKO) mice, generously provided from the laboratory of Khashayarsha Khazaie Ph.D (27), were bred in-house to generate the experimental mice used in this study. We previously demonstrated that cBmalKO mice (experimental group) exhibit mild colon inflammation compared to Ts4-Cre mice (control group) [S]. At 3 weeks of age, male and female Ts4-Cre and cBmalKO mice were weaned and group housed 2 to 5 mice per cage. Mice were maintained in a pathogen-free facility, subjected to a 12/12 hour light/dark cycle, had ad libitum access to standard laboratory rodent chow and water, and were sacrificed by CO_2_ inhalation followed by cardiac puncture at the age of 8 weeks (n = 9-10/genotype/sex). All tissues were harvested between 10 a.m. and 12 p.m.

### Specimen harvesting and preparation

After 2 hours of fasting, body weight (Suppl. Table 1) was measured and blood samples were collected by cardiac puncture. Serum was collected after centrifuging blood samples and stored in −80°C until analysis. Both femurs and L5 vertebra were cleaned of soft tissue and the right femur and L5 vertebra were wrapped in saline-soaked gauze and stored at −20°C and the left femur was fixed and stored in 70% ethanol at room temperature.

### Serum biochemistry

Serum levels of procollagen type 1 N-terminal propeptide (P1NP, Rat/Mouse P1NP EIA, IDS, Gaithersburg, MD) and collagen type 1 C-telopeptide (RatLaps CTX-1 EIA, IDS, Gaithersburg, MD) were evaluated as per the manufacturer’s instructions.

### Microcomputed tomography

Micro-computed tomographic (μCT) imaging was performed on the distal metaphysis and mid-diaphysis of the right femur and the whole L5 vertebra using a high-resolution laboratory imaging system (μCT50, Scanco Medical AG, Brüttisellen, Switzerland) in accordance with the American Society of Bone and Mineral Research (ASBMR) guidelines for the use of μCT in rodents (28). Scans were acquired using a 7.4 μm^3^ isotropic voxel, 70 kVp and 114 μA peak x-ray tube potential and intensity, 300 ms integration time, and were subjected to Gaussian filtration. The distal metaphyseal region for analysis of trabecular bone began 200 μm (27 slices) proximal to the distal growth plate and extended proximally 10% of the femur length, and the trabecular compartment was segmented from the cortical bone by manual contouring. In L5 vertebra, cortical bone was separated from cancellous bone by manual contouring and the region of interest included the region between the end plates. Cortical bone morphology was evaluated in the femoral mid-diaphysis in a region that started at 55% of the bone length proximal to the femoral head and extended 10% of the femur length distally. Thresholds of 350 and 460 mg HA/cm^3^ were used for evaluation of trabecular and cortical bone, respectively. Trabecular bone outcomes included trabecular bone volume fraction (BV/TV, mm^3^/mm^3^), thickness (Tb.Th, mm), and separation (Tb.Sp, mm). Cortical bone outcomes included cortical tissue mineral density (Ct.TMD, mg HA/cm^3^), cortical thickness (Ct.Th, mm), total cross-sectional, cortical bone, and medullary areas (TA, BA, and MA, mm^2^), and the maximum and minimum moments of inertia (I_max_ and I_min_, mm^4^).

### Mechanical testing

The right frozen femurs were thawed and subjected to three-point bending by a materials testing machine (Criterion 43, MTS Systems, Eden Prairie, MN) (29). To determine the stiffness (N/mm) and max load (N), the femur was loaded to failure on the anterior surface at a constant displacement rate of 0.03 mm/sec with the two lower support points spaced 8 mm apart (30). Force-displacement data were acquired at 30 Hz. The frozen L5 vertebrae were thawed and subjected to compression testing at a constant displacement rate of 0.02 mm/sec. The bottom endplate was fixed by cyanoacrylate glue. Force-displacement data were acquired at 30 Hz and stiffness and max load were calculated. Based on the μCT and mechanical testing data, we estimated the cortical bone elastic modulus following methods previously published (29).

### Static and dynamic histomorphometry

Static and dynamic histomorphometric analyses were performed according to the criteria established by the ASBMR (31). Calcein was administered at 2 and 7 days prior to sacrifice. Femurs were dehydrated and embedded in poly methyl methacrylate. 5 μm thick coronal sections were stained with Goldner’s Trichrome for evaluation of static histomorphometric parameters (osteoblast surface/bone surface and osteoclast surface/bone surface) (32) or left unstained for evaluation of the fluorochrome labels. Mineralizing surface per bone surface (MS/BS, %) and mineral apposition rate (MAR, μm/day) were measured on unstained sections to calculate bone formation rate (BFR, μm^3^/μm^2^/day). All measurements were performed using an Osteomeasure image analyzer.

### Statistical analysis

All data were checked for normality, and standard descriptive statistics were calculated. Two-way ANOVA was used to test the main effects of Bmal1 deletion (gene) and sex and their interaction on outcome parameters. Tukey’s HSD post hoc comparisons of means test was used to identify significant differences between groups. Differences were considered significant at *p* < 0.05. Data are reported as mean ± SD.

## Results

Distal femoral metaphyseal trabecular bone mass was reduced by 35% due to Bmal1 deletion in male mice (Figure 1A). This was driven by thinning of trabeculae (−16%) and increased trabecular separation (+13%). L5 vertebral trabecular bone mass was decreased by 21% in male mice due to Bmal1 deletion (Figure 1B). This was primarily driven by thinning of trabeculae by 14% as trabecular separation remained similar. Deletion of Bmal1 in female mice did not lead to alterations in trabecular bone properties of the distal metaphyseal femur or L5 vertebra.

**Figure 1.**
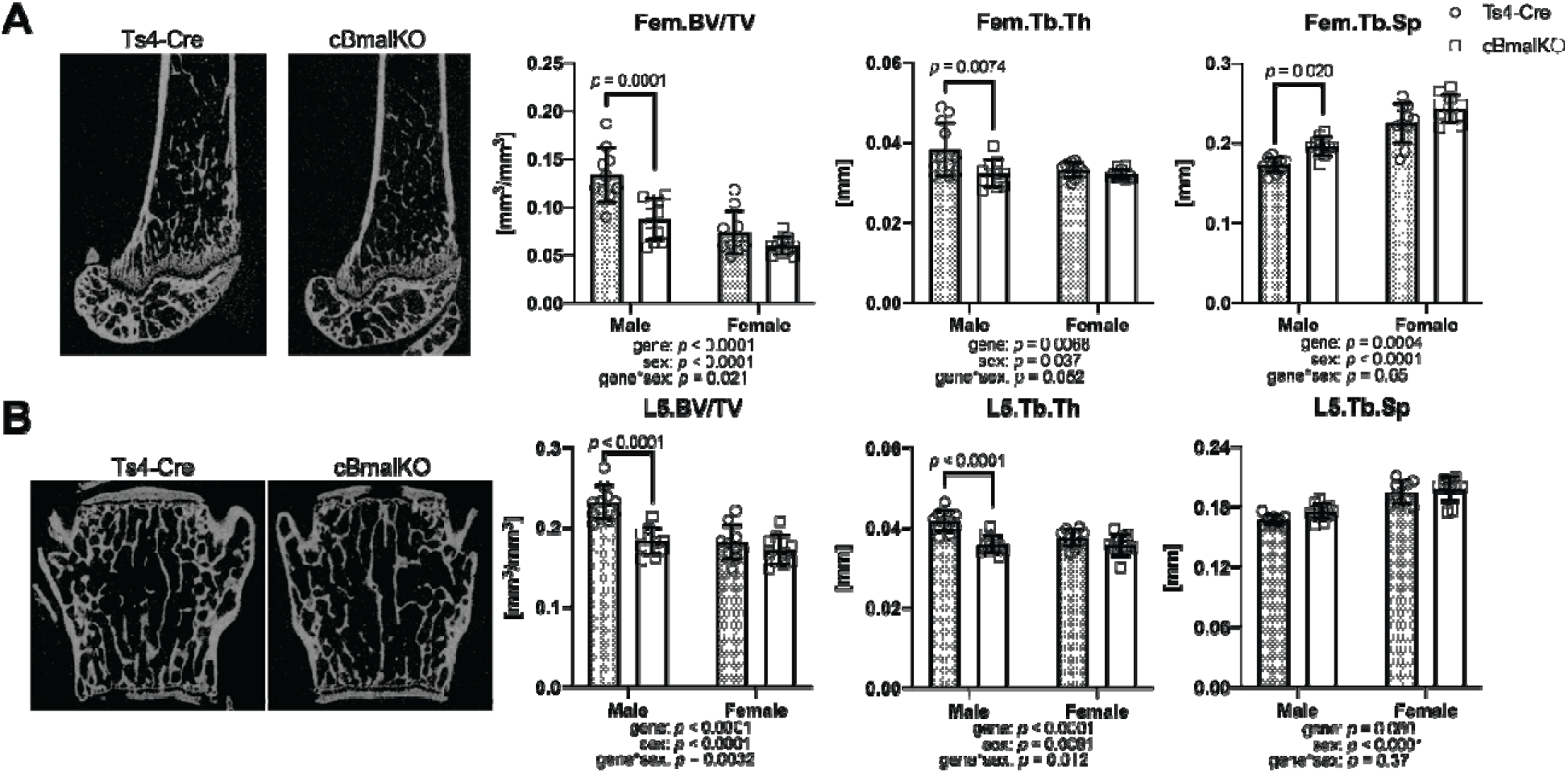
Microarchitecture properties of distal metaphyseal femur (A) and L5 vertebra (B). μCT images from male Ts4-Cre and cBmalKO mice. Fem = femoral; L5 = L5 vertebral; BV/TV = bone volume/total volume; Tb.Th = trabecular thickness; Tb.Sp = trabecular separation. Data presented as Mean ± SD

In cBmalKO male mice, cortical bone was thinner by 13% compared to Ts4-Cre male mice (Table 2). Similarly, total area (−12%), bone area (−16%), maximum moment of inertia (−25%), and minimum moment of inertia (−29%) were lower in cBmalKO male mice compared to Ts4-Cre male mice while medullary area and cortical bone tissue mineral density remained the same. In female mice, Bmal1 deletion led to higher cortical bone tissue mineral density (+7.5%).

**Table 2.**
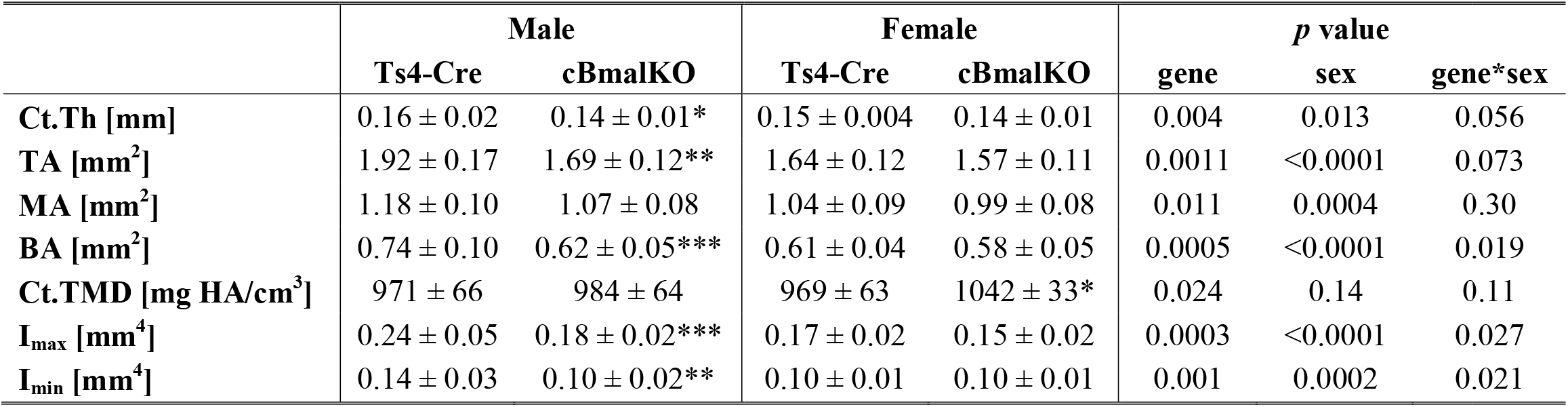
Diaphyseal femoral cortical bone parameters. Ct.Th = cortical thickness; TA = total area; MA = medullary area; BA = cortical bone area; Ct.TMD = cortical bone tissue mineral density; I_max_ = maximum moment of inertia; I_min_ = minimum moment of inertia. Data presented as Mean ± SD; \*\*\**p* < 0.0005; \*\**p* < 0.005; \**p* < 0.05; *vs.* Ts4-Cre within the same sex

Femoral diaphyseal stiffness (−23%) and max load (−26%) were lower in cBmalKO male mice compared to Ts4-Cre male mice (Fig 2). While L5 vertebral stiffness was not altered by the Bmal1 deletion, max load was lower by 26% in cBmalKO male mice. Similar to other skeletal parameters, female cBmalKO mice exhibited comparable biomechanical properties to Ts4-Cre female mice.

**Figure 2.**
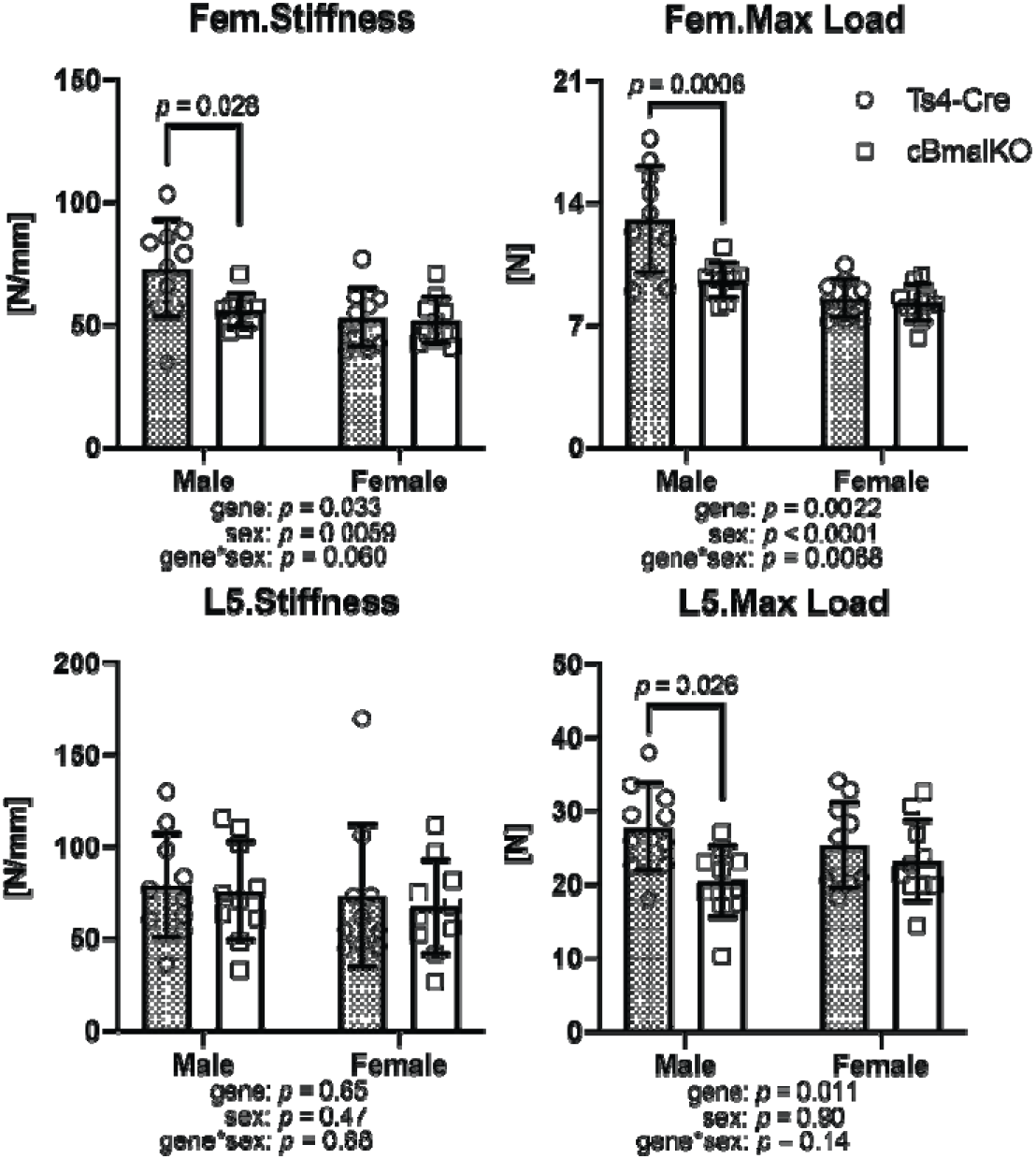
Biomechanical properties of femoral diaphysis and L5 vertebrae in Ts4-Cre and cBmalKO mice. Fem = femoral; L5 = L5 vertebral. Data presented as Mean ± SD

To determine if Bmal1 deletion alters bone resorption and formation, we assessed serum markers of bone turnover by ELISA and static and dynamic histomorphometry of distal femoral trabecular bone. Serum P1NP was 30% lower in cBmalKO mice compared to Ts4-Cre mice in the males, while no differences were observed between the two groups in female mice (Fig 3A). Serum CTX-1 was unaffected by Bmal1 deletion or sex.

**Figure 3.**
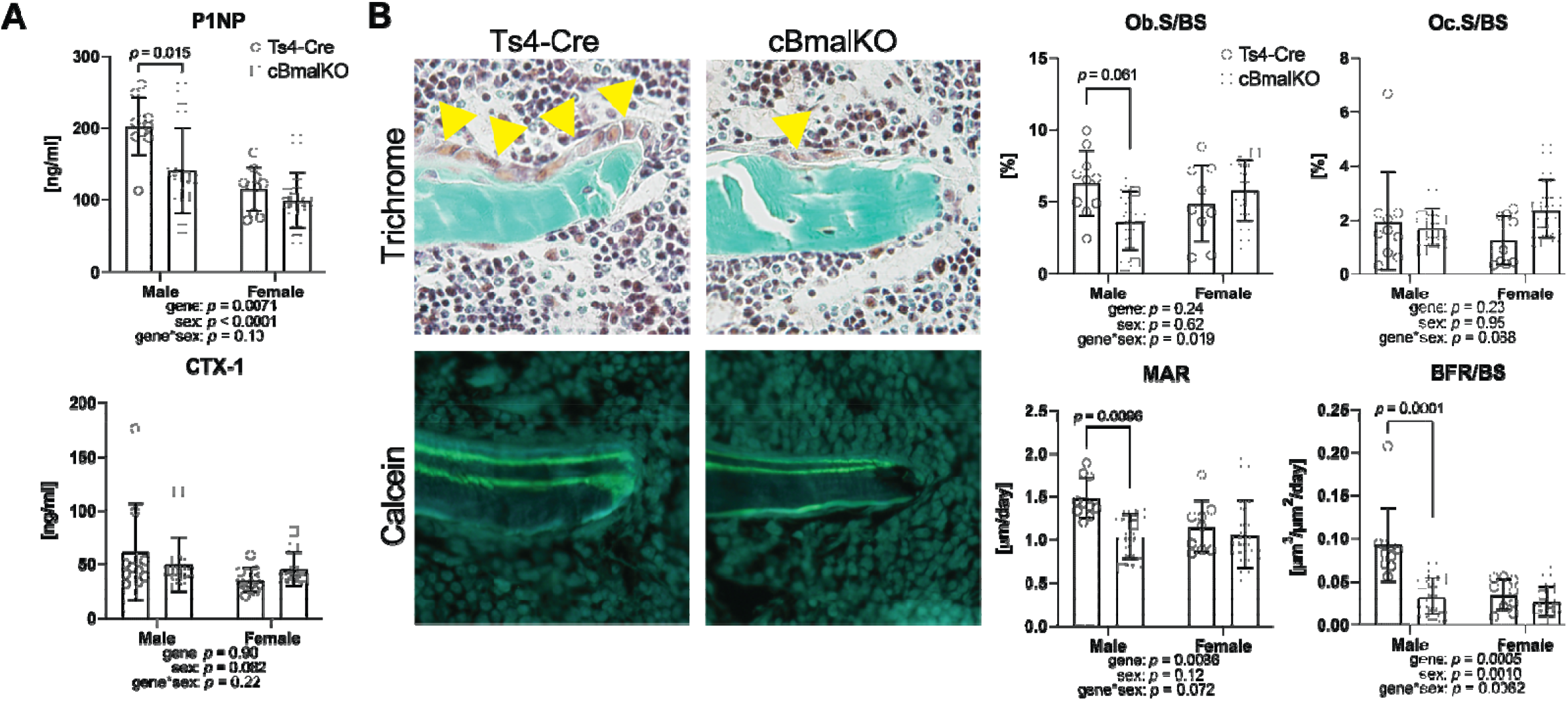
Serum bone remodeling markers (A) and static and dynamic histomorphometry (B). Trichrome and fluorochrome images from male Ts4-Cre and cBmalKO mice. Ob.S/BS = osteoblast surface/bone surface; Oc.S/BS = osteoclast surface/bone surface; MAR = mineral apposition rate; BFR/BS = bone formation rate/bone surface. Data presented as Mean ± SD

Static and dynamic histomorphometric parameters further confirmed that Bmal1 deletion impairs bone formation (Fig 3B). In male cBmalKO mice, Ob.S/BS was lower by 42% compared to Ts4-Cre mice whereas Oc.S/BS was similar. Bmal1 deletion in colon also led to 30% lower MAR and 65% lower BFR/BS in male mice, but not in the females.

## Discussion

This is the first study to demonstrated that colon epithelial cell-specific deletion of Bmal1 leads to skeletal deterioration. The effect includes both cortical and trabecular bone with consequences for biomechanical properties in the appendicular and axial skeleton, but only in male mice. These differences appear to be driven by impaired bone formation in cBmalKO male mice, both evidenced by both the serum bone formation marker P1NP and dynamic histomorphometry.

Our findings are consistent with a previous study where deletion of Bmal1 in cells expressing Villin-Cre was associated with decreased bone (18), but the tissue level mechanism was not consistent (no change in bone resorption vs. elevated resorption in the other study). The differences may be attributed to the specificity of Bmal1 deletion, where Villin-Cre deletes Bmal1 in both the small and large intestine and the kidney, whereas Ts4-Cre used in our studies only deletes Bmal1 in the large intestine and the very distal ileum (19, 33). The age at which the skeletal phenotyping was performed may also explain the differences observed (8 weeks old in our study *vs.* 16 weeks old).

The age (8 weeks old) at which we assessed the skeleton of Ts4-Cre and cBmalKO mice corresponds to the age when mice reach their peak trabecular bone mass in both the appendicular and axial bone (34). In the femoral diaphysis, the cross-sectional area and cortical bone thickness continue to increase beyond 8 weeks (35). Our findings suggest that in male mice, the acquisition of peak trabecular bone mass is impaired by colon-specific Bmal1 deletion. The architectural and functional deficits in the femoral diaphysis may continue to persist as the mice age, but further study is needed to confirm that. Whether female mice will continue to be protected from skeletal deterioration as they age is unclear.

Our study surprisingly revealed that female mice were protected from structural and functional skeletal deficits due to Bmal1 deletion in colon epithelial cells. Studies that examine whether the gut-bone axis exhibits sexual dimorphism are limited, but a previous study demonstrated that female mice were protected from increased gut permeability from antibiotics and subsequent deteriorations in the skeleton (36). Also, ulcerative colitis patients showed decrease in bone mineral density that was more significant in male than in female patients (37). Since studies have suggested that estradiol has protective effects against metabolic disorders such as obesity, osteoporosis, and diabetes (38–40), female mice in our study may similarly be protected from colon-specific Bmal1 deletion-mediated skeletal deterioration due to higher levels of estradiol (41, 42).

We acknowledge the limitation of our studies where the underlying molecular mechanisms of colon Bmal1 deletion-mediated skeletal deterioration were not examined, although we did identify one of the tissue level mechanisms (suppressed bone formation). While the mild inflammation observed in colon in this model [S] may suggest osteoclast-mediated bone resorption as shown in other studies (43–45), our studies of serum bone remodeling markers and dynamic histomorphometry suggest that Bmal1 deletion is likely inhibiting factors in the colon that promote bone formation. Also, although we observed mild colon inflammation, cBmalKO mice did not exhibit increased gut leakiness compared to Ts4-Cre mice [S], suggesting that other gut-derived factors are likely regulating skeletal homeostasis. Gut microbiome and gut-derived hormones are known to regulate osteoblast function and thereby bone formation (46, 47). The circadian rhythm of gut-derived plasma short chain fatty acids (SCFAs) has been shown to be disrupted among shift workers (48), and along with a previous study that shows the importance of SCFA butyrate to bone formation (49), the skeletal pathologies seen in our studies may be attributed to the disruption of gut microbiota due to a disruption in circadian signaling in gut.

In conclusion, our study demonstrates that colon epithelial cell-specific deletion of Bmal1 leads to trabecular and cortical bone loss in male mice, whereas female mice are unaffected. This suggests that strategies that maintain the circadian rhythm in colon may prevent subsequent skeletal deterioration observed in sleep-deprived individuals.

## Author Contributions

Study design: FCK, CF, AK, GRS, DRS. Study conduct and data collection: FCK, SBJ, BMW, AA, NP, RM, SW, MS. Data analysis: FCK, DRS. Drafting and revising manuscript content: all authors. Approving final and submitted version of manuscript: all authors. DRS takes responsibility for the integrity of the data analysis.

## Acknowledgments

This study was supported by NIH R21AR075l30 (DRS), T32AR073157 (DRS), R24AA026801 (AK), K01AR077679 (FCK), and Pfizer Competitive Grant Program: Inflammatory Bowel Disease 2019 (GRS). The content is solely the responsibility of the authors and does not necessarily represent the official views of the National Institutes of Health. AK would also like to acknowledge philanthropy funding from The Johnson Family, Mrs. Barbara and Mr. Larry Field, Mrs. Ellen and Mr. Philip Glass, Mrs. Marcia and Mr. Silas Keehn and the Sklar Family. We thank Dr. Khashayarsha Khazaie for his scientific support for this project. Rush University Medical Center MicroCT/Histology Core provided experimental support.

